# A Mathematical Methodology to Predict Phenotype from Genotype

**DOI:** 10.1101/2025.03.17.643655

**Authors:** Jeonghoon Lee, Kyu Yeong Choi

**Author notes:** Corresponding Author : (E-mail) (Jeonghoon Lee).

## Abstract

From a mathematical perspective, a genome can be viewed as categorical data whose elements include genetic variants in SNPs. Then, the problem of predicting phenotype from genotype can be reduced to the problem of classifying categorical data. We will define a metric function at each SNPs position of an individual such that the arithmetic mean of a specific phenotype group is always less than or equal to that of the comparison group. Then, in light of a mathematical principle, calculating the sum of the distances(metrics) of some SNPs positions could clearly classify the two groups, when difficult to classify them with one specific position.

Using this methodology, we found that even small differences in each SNPs combined could classify phenotypes ; combining 51 SNPs positions in autosomes made possible to distinguish males from females with great accuracy. We also found that East Asian people could be distinguished from other people with 100% accuracy, so we obtained a mathematical definition of East Asian people.

The mathematical methodology classifies phenotypes with collecting small differences of genotypes throughout the entire genome, which could be useful for predicting diseases affected by the genome and be useful in pharmacogenomics.

## 1. Introduction

Single nucleotide polymorphisms(SNPs) are associated with phenotypic expressions(Shastry).^1^ Genome-wide association studies(GWAS) are to identify associations of genotypes with phenotypes commonly studying on genetic variants in SNPs(Uffelmann *et al*).^2^ The function of SNPs is still being studied, but not much is known yet(Visscher, P. M. *et al*.^3^ Khera, A. V. *et al*.^4^ Peterson, R. E. *et al*.^5^ Lello, L., *et al*.^6^ Watanabe, K., *et al*.^7^).

From a mathematical perspective, a genome can be viewed as categorical data, and genetic variants in SNPs can be viewed as elements that constitute the data. Then, the problem of predicting phenotype from genotype can be reduced to the problem of classifying categorical data. In this case, even if the function of the variants in SNPs is not known exactly, the phenotype can be predicted from the genotype.

Since a phenotype is the result of a genotype as well as of environmental factors(Taylor *et al*),^8^ it can be assumed that a structural difference in genotype exists when comparing the group of people with a specific phenotype and the comparison group(Amos, W.^9^ Zhanshan Ma., *et al*^10^). The difference in the probability distribution of nucleobases(or genotypes) at the position of a specific SNPs can be used as an indicator to capture structural differences in genomes between groups.

In the case of Mendelian trait, the probability distributions of the genotypes of the dominant group are different from that of the recessive group. In the dominant group there are people with genotype RR or Rr, and in the recessive group only with the genotype rr. If the probabilities of such people within the dominant group are *α* and 1-*α*, respectively, the dominant group’s probability distribution (RR, Rr, rr) = (*α*, 1-*α*, 0) and, the recessive group’s (RR, Rr, rr) = (0, 0, 1), where R is the dominant allele, r the recessive allele, (RR, Rr, rr) the collection of the genotypes, and (*α*,1-*α*,0), (0,0,1) the collection of the probability of each genotype.

In the case of non-Mendelian trait, two or more genes(or SNPs) are involved in the phenotype(Susan Strome., *et al*.),^11^ and this can be expected to appear as differences in the probability distribution of genotypes between phenotype groups at several SNP positions. So it can be assumed as follows:

“At one or more positions of SNPs exists the difference of the probability distribution of nucleobases(or genotypes) between of a group with a specific phenotype and of a comparison group.” (Assumption)

In order to predict phenotypes from genotypes, this paper introduces a mathematical methodology to measure genetic distance between an individual and a group with a specific phenotype under the assumption above. The methodology distinguishes phenotypes by accumulating small differences of the probability distribution of genotypes throughout the entire SNPs.

We will define a metric function at each SNPs position of individual such that the arithmetic mean of a specific phenotype group is always less than or equal to that of the comparison group. Then, we can use the implication that if *N* IID(independent and identically distributed) random variables are summed, the mean becomes *N* times, but the standard deviation becomes 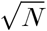 times (Jacobs, K.).^12^ For example, there is not much difference between normal distributions *N*(0.5, 0.1^2^) and *N*(0.55, 0.1^2^) as seen in Figure 1, but *N*(50 1^2^) and *N*(55 1^2^) in Figure 2, summed by 100 IID random variables respectively, have a clear difference, making it easy to classify members of the two groups. That is, calculating the sum of the metrics(distances) of several SNPs positions could clearly classify the members of the two groups, even when difficult to classify with one SNPs position.

**Figure 1.**
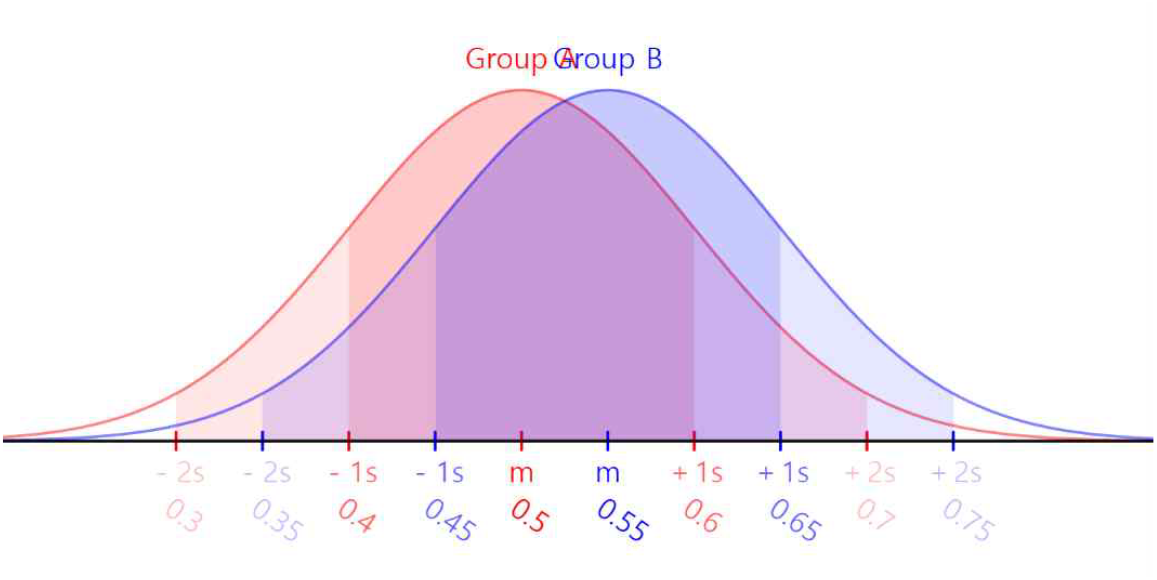
The classification of *N*(*0*.5 *0*.1^2^) and *N*(0.55 0.1^2^). Group A is *N*(0.5, 0.1^2^), Group B *N*(0.55 0.1^2^) (drawn using Javalab, https://javalab.org)

**Figure 2.**
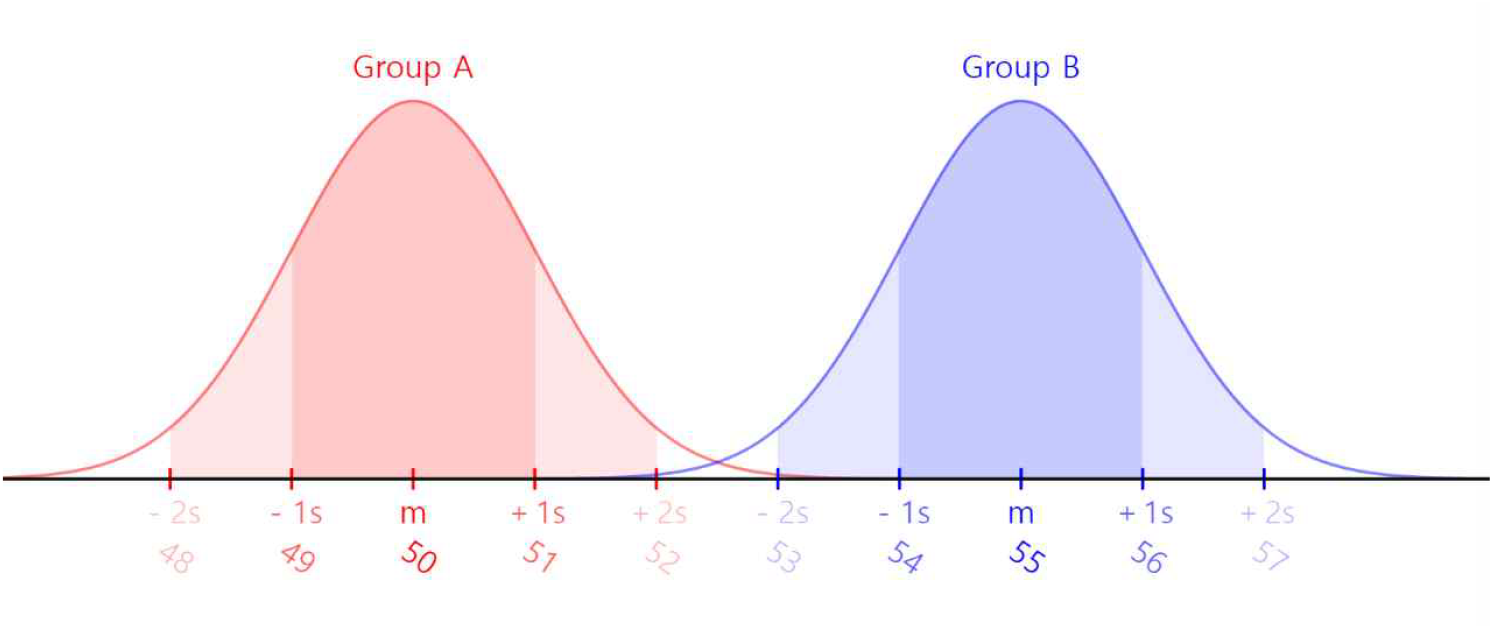
The classification of *N*(50, 1^2^) and *N*(55 1^2^). Group A is *N*(50, 1^2^), Group B *N*(55 1^2^) (drawn using Javalab, https://javalab.org)

The methodology will show that males can be distinguishable from females even excluding sex chromosomes, which implies that between males and females there are genetic differences in autosomes. We also found that East Asian people could be distinguished from other people with 100% accuracy, so we obtained a mathematical definition of East Asian people..

## 2. Categorical distance

Categorical distance measures the distance between an individual and a group with a specific phenotype using the probability distributions of bases(or genotypes), and through this, it is possible to determine whether an individual is included in the group or not.

### Category of genotype

Category of genotype denotes a type of possible genotypes at specific SNPs position, such as reference, alternative 1, alternative 2, etc, and for autosomes, it refers to a collection of possible pairs, such as (reference, reference), (reference, alternative 1), (alternative 1, reference), (alternative 1, alternative 1), etc. Since the two chromosomes of autosome can not be distinguishable by one SNPs position, (alternative 1, reference) is the same as (reference, alternative 1). Reference and alternative could be a base, a plurality of bases, or an indel.

According to GRCh38 version of ‘1000 Genomes Project’, at the position 18212123 of chromosome 13, the reference is A and alternative 1 is G. There are three categories : 0|0, 0|1(=1|0), and 1|1. That is, category 1, category 2, and category 3. (where 0|0=(reference, reference), 0|1=(reference, alternative 1), 1|1 = (alternative 1, alternative 1))

If there are AC, C, CT of three genotypes, the reference is AC, alternative 1 is C, alternative 2 is CT, and there are six possible categories : 0|0, 0|1(=1|0), 1|1, 0|2 (=2|0), 1|2(=2|1), and 2|2.

### Definition of categorical distance

The categorical distance of an individual *x* to group X with a specific phenotype at the position *i* is defined as:

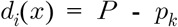

(where, at the position *i, p*_*j*_ is probability of category j, the probability distribution of genotypes of group X is (*p*_*1*_,*p*_2_,…,*p*_*n*_), *P*=max(*p*_1_,*p*_2_,…,*p*_*n*_), and the genotype of *x* is ‘category *k*’)

If at the position 100, (0|0, 0|1, 1|1) = (0.81, 0.18, 0.01), and the genotype of *x* is 0|1, then *d*_100_(*x*) = *P* - *p*_*k*_ = max(0.81, 0.18, 0.01) – *p*_2_ = 0.81 − 0.18 = 0.63.

Categorical distance can be viewed as a measure of how commonly a genotype at a specific position of an individual appears in that group. It changes a genotype from “a letter” to a numerical value, and makes it possible to calculate the mean value of groups.

### Mean of categorical distances

At the position *i*, calculating the arithmetic mean of the categorical distances to group X for all elements(individuals) of group X is as follows:

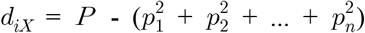

At the position *i*, calculating the arithmetic mean of the categorical distances to group X for all elements(individuals) of group Y is as follows:

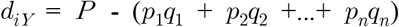

(where, at the position *i*, the probability distribution of genotypes of group X is (*p*_1_,*p*_2_,…,*p*_*n*_), that of group Y is (*q*_1_,*q*_2_,…, *q*_*n*_), and *P*=max(*p*_1_,*p*_2_,…,*p*_*n*_))

### Collective distance

We assumed that the probability distribution of genotypes be different at some SNPs positions between the group X with a specific phenotype and the comparison group Y. This means (*p*_1_,*p*_2_,…,*p*_*n*_) ≠ (*q*_1_,*q*_*2*_,…,*q*_*n*_) for some SNPs positions.

The mean of categorical distance suggests a method of finding such SNPs positions that the probability distribution of genotypes is different. If *d*_*i X*_ ≠ *d*_*i Y*_ at the position *i*, then (*p*_1_,*p*_2_,…,*p*_*n*_) ≠ (*q*_1_,*q*_2_,…,*q*_*n*_) at that position. Thus, it is to find the positions where *D*_*i XY*_ ≡ *d*_*i X*_ - *d*_*i Y*_ ≠ 0. It can be estimated that the larger |*D*_*i XY*_ |, the greater the difference in probability distributions.

In general, it is likely *d*_*i X*_ ≤ *d*_*i Y*_, but sometimes it could be *d*_*i X*_ > *d*_*i Y*_. For example, if for X, (0|0, 0|1, 1|1)=(0.81, 0.18, 0.01) and for Y, (0|0, 0|1, 1|1) = (1, 0, 0) then 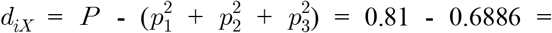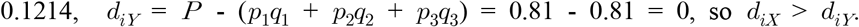.

To modify categorical distance, define collective distance *dS*_*i*_ as follows :

At the position *i*, if *D*_*i XY*_ ≤ 0, then *dS*_*i*_ = *d*_*i*_, otherwise *dS*_*i*_ = 1 - *d*_*i*_

(where *d*_*i*_ is categorical distance)

Thus, *dS*_*i X*_ ≤ *dS*_*i Y*_, at all SNPs positions.

(where *dS*_*i X*_ and *dS*_*i Y*_ are the arithmetic mean of the collective distances of X and Y respectively)

The collective distance *dS*_*i*_ is a metric function defined at each SNPs position of an individual such that the arithmetic mean of a specific phenotype group is always less than or equal to that of the comparison group. In light of the mathematical principle above, calculating the sum of the collective distances of some positions could clearly classify the members of the two groups, even when difficult to classify with one specific position.

It is desirable that the positions are selected where |*D*_*i XY*_ | is large. Because the larger |*D*_*i XY*_ |, the greater the difference in probability distributions so that we could ignore statistical noise.

### Classifying groups

Now let *Q*={*i*_1_,*i*_2_,…,*i*_*m*_} be the selected positions, and define

*dT*_*Q*_, the total collective distance for *Q*, as 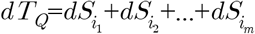.

If an threshold *Δ* is decided, it is possible to determine whether an individual *z* is included in X or not. If *dT*_*Q*_(*z*) ≤ *Δ*, then *z* ∈X.

## 3 Result 1 : Classifying human sexes only with autosomes

### The used data and samples

Among the ‘1000 Genomes Project’ data(The 1000 Genomes Project Consortium),^13^ the GRCh38 version was used. The information of samples of the GRCh38 version data is found at https://www.internationalgenome.org/data-portal/sample, which includes the information of sex. The words of ‘sex’, ‘male’, and ‘female’ are the categories the data adopt.

300 males and 300 females in front based on the list order were excluded for the test and verification. These samples were not used for finding SNPs positions. Using the samples not excluded, the group X(952 males) and the group Y(996 females) were made.

### Finding SNPs positions

For 952 males(X) and 996 females(Y), we evaluated |*D*_*i XY*_ | of each SNPs position from chromosome 1 to chromosome 22. Then, we found the positions P1 where |*D*_*i XY*_ | > 0.6, and the positions P2 where 0.07 < |*D*_*i XY*_ | < 0.21 (no positions found with 0.21 ≤ |*D*_*i XY*_ | ≤ 0.6). The positions found are listed on the Supplementary Materials.

### Calculating total collective distance

For the excluded 300 males and 300 females who were excluded for the test and verification, we calculated the total collective distance of each individual for the positions(P1) where |*D*_*i XY*_ | > 0.6, and for the positions(P2) where 0.07 < |*D*_*i XY*_ | < 0.21 respectively. The results are on Supplementary Materials.

In the autosomes, there was almost no difference between males and females. However, there were some differences in approximately 60 SNPs positions.

### Analysis of SNPs positions(P1) where |*D*_*i XY*_ | > 0.6

There were 6 positions including rs1169137959 of chromosome 13. Other 5 positions were on chromosome 17; rs1304242437, rs1358569312, rs1412643511, rs1472916410, rs1389701773.(dbSNP identifier based). The mean of the total collective distance of 300 males was 0.267; the standard deviation was 0.476. The mean of the total collective distance of 300 females was 5.224; the standard deviation was 0.178. Males were distinguishable from females using these SNPs positions. With the threshold 3, the 299 males and the 0 female were below the threshold(the accuracy : 99.8%.) (Figure.3)

**Figure 3.**
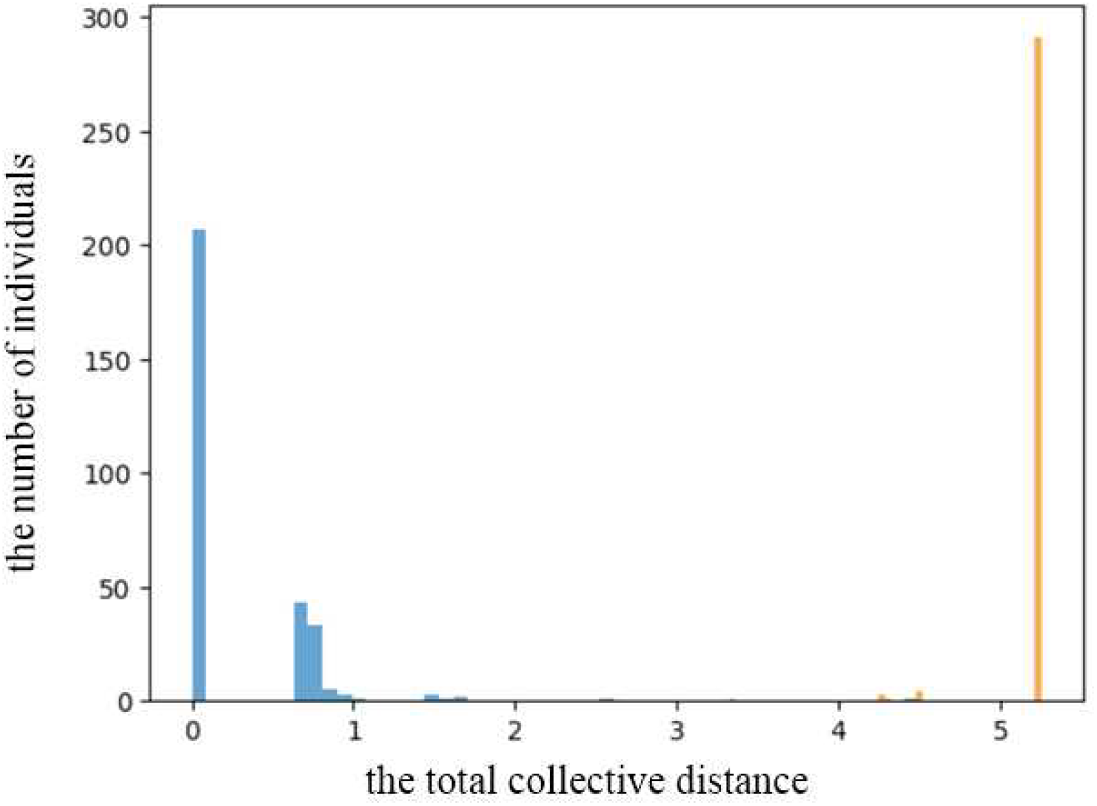
The classification of human sexes by using the positions of large differences in autosomes. The horizontal axis is the total collective distances for 6 SNPs positions (|*D*_*i XY*_ | > 0.6), and the vertical axis is the number of individuals included in the range. Blue is males, orange is females. If the threshold *Δ*=3, the accuracy is 99.8%. Python with “plt.hist(males/females, bins=50, density=False, alpha=0.7, histtype=‘bar’)” was used for the figure

### Analysis of SNPs positions(P2) where 0.07 < |*D*_*i XY*_ | < 0.21

There were 51 positions in autosomes with 0.07 < |*D*_*i*_*XY*| < 0.21. The mean of the total collective distance of 300 males was 27.586; the standard deviation was 2.132. The mean of the total collective distance of 300 females was 32.643; the standard deviation was 1.214.

Although the difference in each position was not significant, combining 51 positions made possible to classify human sexes with great accuracy. With the threshold 30.5, the 285 males and the 13 females were below the threshold. (the accuracy : 95.3%) It was showed that small differences in SNPs could classify phenotypes as a whole. (Figure.4)

**Figure 4.**
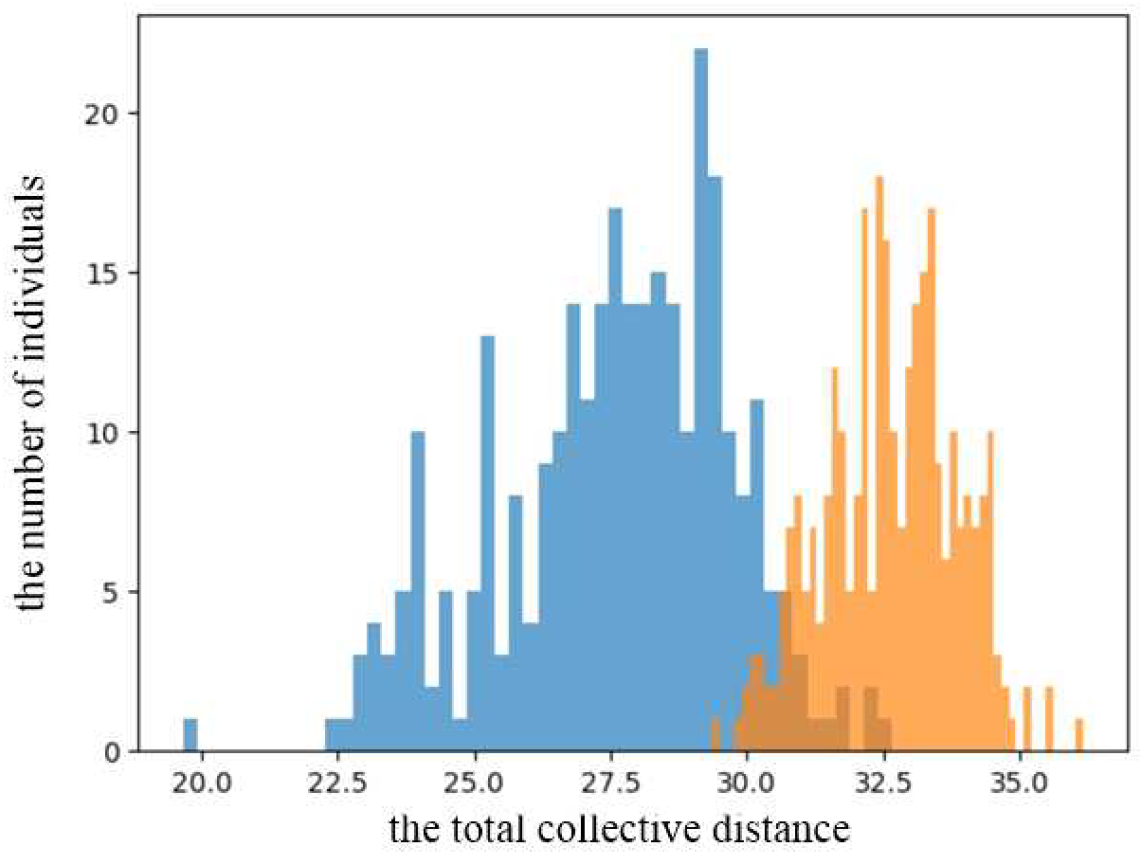
The classification of human sexes by using the positions of small differences in autosomes. The horizontal axis is the total collective distances for 51 SNPs positions(0.07 < |*D*_*i XY*_ | < 0.21), and the vertical axis is the number of individuals included in the range. Blue is males, orange is females. If the threshold *Δ*=30.5, the accuracy is 95.3%. Python with “plt.hist(males/females, bins=50, density=False, alpha=0.7, histtype=‘bar’)” was used for the figure.

### Statistical information

We use the Welch’s t-test since the variances of two groups are not equal, and calculate p-value from t-test results.

The standard error of the mean (SE) is as follows:

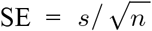, where *s* is the standard deviation of the sample, and *n* is the sample size.

The t-value(test statistic) is as follows:

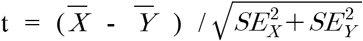, where 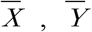 are the sample mean of X, Y respectively, and *SC*_*X*_, *SC*_*Y*_ are the SE of X, Y respectively.

The degrees of freedom for Welch’s t-test are as follows:

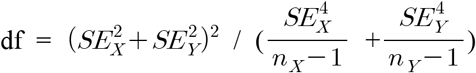, where *n*_*X*_, *n*_*Y*_ are the sample size of X, Y respectively.

Null Hypothesis (H0) and Alternative Hypothesis (H1) are as follows:

H0 : *μ*_*X*_ *= μ*_*Y*_, H1 : *μ*_*X*_ ^3^≠ *μ*_*Y*_, where *u*_*X*_, *u*_*Y*_ are the mean of X, Y respectively.

For the positions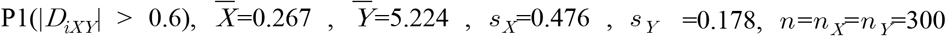, then t ≒-177, df ≒ 353 The critical t-value(two-tails) in the t-table is 7.0413 which is corresponding to significance level α=1×*l0*^*—*11^and degrees of freedom df=353. Since | t | =177 > 7.0413, then the null hypothesis is rejected. In addition, when t=177, df=353, p-value(two-tails) is less than 1.1877×*l0*^*—*15^; it is too small to calculate for a computer.

For the positions P2 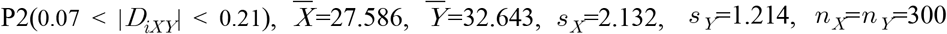 then t ≒-35.76, df≒ 472.9. The critical t-value(two-tails) in the t-table is 6.9808 corresponding to significance level, α =1×10^*—* 11^ and degrees of freedom df=472. Since | t | =35.76 > 6.9808, then the null hypothesis is rejected. In addition, when t=35.76, df=472, p-value(two-tails) is less than 5.8127×10^*—*18^; it is too small to calculate for a computer.

## 4 Result 2 : Distinguishing East Asian people from other people

### Samples

The information of samples of the GRCh38 version data was found at https://www.internationalgenome.org/data-portal/sample, which includes the information of superpopulation. ‘EAS’, ‘AFR’, ‘AMR’, ‘EUR’, and ‘SAS’ are the categories of superpopulation the data adopt.

There were 515 East Asian people(EAS), and 2032 other people(AFR, AMR, EUR, and SAS)(one sample of double superpopulations was excluded)

### Finding SNPs positions

Using 515 East Asian people(X) and 2032 other people(Y), we evaluated |*D*_*i XY*_ | of each SNPs position from chromosome 1 to chromosome 22. Then, we found the positions where |*D*_*i XY*_ | > 0.6.

### Calculating total collective distance

For 515 East Asian people(X) and 2032 other people(Y), we calculated the total collective distance of each individual for the positions where |*D*_*i XY*_ | > 0.6.

### Analysis of SNPs positions where |*D*_*i XY*_ | > 0.6

There were 4,888 positions from chromosome 1 to chromosome 22 where |*D*_*i XY*_ | > 0.6. The mean of the total collective distance of East Asian people(EAS) was 199.839; the standard deviation was 100.118. The mean of the total collective distance of other people was 3321.674; the standard deviation was 594.622. All the East Asian people were less than 730, and all other people were larger than 1324. So if the threshold is set to 1000, the accuracy becomes 100%. (figure 5)

**Figure 5.**
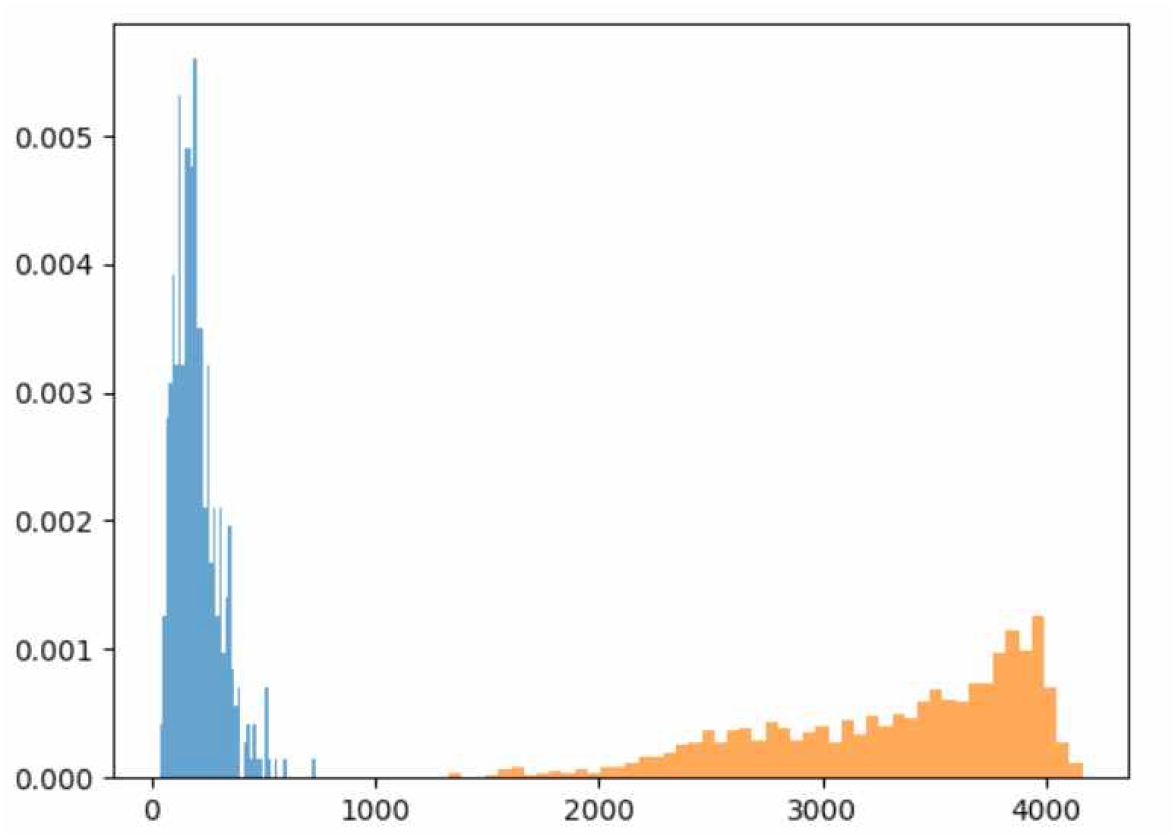
East Asian people(EAS) and other people(AFR, AMR, EUR, and SAS). The horizontal axis is the total collective distances for 4,888 SNPs positions(|*D*_*i XY*_ | > 0.6), and the vertical axis is ‘the number of individuals(converted to percentage)’ included in the range. Blue is East Asian people(EAS), orange is other people. If the threshold *Δ*=1000, the accuracy is 100%. Python with “plt.hist(men/women, bins=50, density=True, alpha=0.7, histtype=‘bar’)” was used for the figure.

### Definition of East Asian people(EAS)

Let us define East Asian people as EAS which is the category of superpopulation the data adopt. Since we can distinguish East Asian people from other people with 100% accuracy, we are able to define East Asian people as follows : “A member of East Asian people(EAS) denotes a person whose total collective distance for 4,888 SNPs positions(|*Di*_*XY*_| > 0.6) is less than 1000.”

## 5 Discussion

We could consider drug response or disease susceptibility as a phenotype. The mathematical methodology of this paper to predict phenotypes from genotypes could be useful for predicting diseases affected by the genome and useful in pharmacogenomics. For example, it could help predict Alzheimer’s disease or identify people who respond to cancer immunotherapy in advance.

From a theoretical perspective, let X, Y be a phenotype group and its comparison group respectively, for the SNPs positions that classify the two groups, we can compute the average of the total collective distances of the other phenotype group Z, which allows us to determine whether Z is closer to X or Y. In other words, we can measure the similarity between phenotypes. We can also specify the SNPs positions that cause such similarity. If we accumulate statistics, we could identify the functions of SNPs or set of SNPs and understand genome better.

In addition, rs1169137959 of chromosome 13 was discovered, which shows that the egg cell’s genotype is an important factor in determining human sex; the genotype of most men(87.86%) is (A, A), and that of most women(97.69%) is (A, G). (Appendix)

## Appendix

### Egg Cell’s Genotype Determining Human Sex

#### The finding facts

The GRCh38 version was for 1,252 males and 1,296 females, and at the position of rs1169137959, the reference was A(adenine) and the alternative was G(guanine).

For males, there were 1,100(87.86%) with genotype (A, A), 152(12.14%) with (A, G).(none with (G, G))

For females, there were 30(2.31%) with genotype (A, A), 1,266(97.69%) with (A, G).(none with (G, G))

#### Demonstration

Statistics show that the genotype of most males is (A, A), and that of most women is (A, G).

After meiosis, if [x, A], [y, A] of the sperm and [X, A], [X, G] of the egg combine to show the same survival probability, then in the next generation, immediately [xX, (A,A)] and [xX, (A,G)] survive in similar numbers, and the probabilities of genotype (A, A) and (A, G) in females will be similar, where [x, A] is a sperm with chromosome X and genotype A at rs1169137959, and [xX, (A, A)] is a female with genotype (A, A), etc.

However, the statistics show a large difference between females with (A, A)(2.31%) and females with (A, G)(97.69%). This shows that there is a great difference in survival probability between [x, A]&[X, A] combination and [x, A]&[X, G] combination. That implies the survival probability of [x, A]&[X, G] combination is much greater. Here ‘survival probability’ refers to the probability of passing the genome to the next generation.

For the same reason, the survival probability of the combination of [y, A]&[X, A] is much greater than that of the combination of [y, A]&[X, G], where [y, A] is a sperm with chromosome Y and genotype A at rs1169137959, and [X, A] is a egg cell with genotype A, etc.

In conclusion, [X, A] type eggs mainly combine with [y, A] type sperms, and [X, G] type eggs mainly combine with [x, A] type sperms, and other combinations have a low probability of survival. That is, most males are the result of combining with [X, A] type eggs, and most females are the result of combining with [X, G] type eggs.

Therefore, an important factor in determining human sex is the genotype of the egg cell.

#### List of Supplementary Materials (classifying human sexes)

SNPs positions where |*D*_*i XY*_ | > 0.6

SNPs positions where 0.07 < |*D*_*i XY*_ | < 0.21

The total collective distances for 6 SNPs (|*D*_*i XY*_ | > 0.6)

The total collective distances for 51 SNPs (0.07 < |*D*_*i XY*_ | < 0.21)

## Data availability

The GRCh38 version of “1000 Genomes Project” data is available: https://www.internationalgenome.org

## Acknowledgements

We would like to express our gratitude and appreciation to Dr. Sangnam Nam, Dr. Kyoung-Seok Ryu and Mr. Daeman Shin for valuable advice.

## Author contributions

J.H.L. conceived the idea, designed the study, performed the data analysis, and wrote the manuscript draf ; K.Y.C. interpreted the results and revised the manuscript.

## Materials & Correspondence

Correspondence and requests for materials should be addressed to Jeonghoon Lee.

