## Supplementary Materials for "A Mathematical Methodology to Predict Phenotype from Genotype"

### List of Supplementary Materials

#### (classifying human sexes)

1.1. SNPs positions where $|D_{iXY}| > 0.6$

1.2. SNPs positions where $0.07 \leq |D_{iXY}| \leq 0.21$

1.3. The total collective distances for 6 SNPs ($|D_{iXY}| > 0.6$)

1.4. The total collective distances for 51 SNPs ($0.07 \leq |D_{iXY}| \leq 0.21$)

#### 1.1. SNPs positions where $|D_{iXY}| > 0.6$

(position number in GRCh38, $|D_{iXY}|$ ) : dbSNP ID

chr 13

('18212123', 0.6444051752522412) : rs1169137959

chr 17

('26867060', 0.9637934550148906) : rs1304242437

('26867146', 0.9250370594729741) : rs1358569312

('26866764', 0.8976834266662015) : rs1412643511

('26866745', 0.822476758244049) : rs1472916410

('26867019', 0.6064387484263727) : rs1389701773

#### 1.2. SNPs positions where $0.07 \leq |D_{iXY}| \leq 0.21$

(position number in GRCh38, $|D_{iXY}|$ )

chr 1

('242917670', 0.20716295908866728), ('242886341', 0.11797898642093263), ('242887048', 0.11397115543595182), ('242911992', 0.10097135948638841), ('202378896', 0.07369052333903087)

chr 2

('88773387', 0.12181343125485489), ('88773398', 0.12181343125485489), ('95938800', 0.11903290728055926), ('87883796', 0.1132570941204264), ('97168870', 0.10380622837370235), ('87905256', 0.10024715059659584), ('87905255', 0.1000480277156931), ('87882867', 0.07631820446108994), ('87870325', 0.07146208057992864)

chr 3

('16438051', 0.11841176052277752), ('16544144', 0.09581148192799444), ('16373342', 0.09115776005653853), ('16551924', 0.08878082593774572)

chr 5

('10346475', 0.09500626083176)

chr 9

('31120122', 0.11553006213116007)

chr 10

('38788107', 0.11762254512010323), ('38879848', 0.08158431693017393), ('38879852', 0.08158431693017393), ('38879885', 0.07604762315982383)

chr 13

('19389753', 0.1391035950170288), ('18211952', 0.08587694468290497), ('18212061', 0.07842812000774801), ('19394927', 0.07470229409978024)

chr 16

('33741748', 0.07981159331202359)

chr 17

('21863032', 0.12462705670503493), ('21863043', 0.12450347786173299),

('26866572', 0.12274027257962017), ('21881265', 0.1212904279358803),

('26863513', 0.12078728550243634), ('26854337', 0.1129062539260921),

('26854338', 0.1129062539260921), ('26854346', 0.1129062539260921),

('26854349', 0.1129062539260921), ('26854355', 0.1129062539260921),

('21903582', 0.10617408728197164), ('21903588', 0.10617408728197164),

('21903589', 0.10617408728197164), ('21929921', 0.10408699269927668),

('21933986', 0.08478170680036723), ('21893345', 0.08072059440360124),

('21863754', 0.07334183673469397)

chr 21

('10735603', 0.09106878045335787), ('10735583', 0.08236309229574179),

('10735592', 0.08236309229574179), ('10735593', 0.08236309229574179),

('10735586', 0.08077269710151957)

**1.3. The total collective distances for 6 SNPs($|D_{iXY}|$ > 0.6)**

**(men) : sample 1(HG00096), sample 2, sample 3, ... ,sample 300(HG01840)**

**the total collective distance of sample 1(0.0), the total collective distance of sample 2, the total collective distance of sample 3, ..., the total collective distance of sample 300(0.7100840336134453)**

'HG00096', 'HG00101', 'HG00103', 'HG00105', 'HG00107', 'HG00108', 'HG00109', 'HG00112', 'HG00113', 'HG00114', 'HG00115', 'HG00116', 'HG00117', 'HG00119', 'HG00126', 'HG00129', 'HG00131', 'HG00136', 'HG00138', 'HG00139', 'HG00140', 'HG00141', 'HG00142', 'HG00143', 'HG00145', 'HG00148', 'HG00149', 'HG00151', 'HG00152', 'HG00155', 'HG00156', 'HG00157', 'HG00159', 'HG00160', 'HG00181', 'HG00182', 'HG00183', 'HG00185', 'HG00186', 'HG00187', 'HG00188', 'HG00189', 'HG00190', 'HG00234', 'HG00242', 'HG00243', 'HG00244', 'HG00246', 'HG00251', 'HG00252', 'HG00256', 'HG00260', 'HG00264', 'HG00265', 'HG00267', 'HG00271', 'HG00273', 'HG00277', 'HG00278', 'HG00280', 'HG00284', 'HG00290', 'HG00303', 'HG00308', 'HG00310', 'HG00311', 'HG00312', 'HG00321', 'HG00325', 'HG00329', 'HG00335', 'HG00336', 'HG00338', 'HG00341', 'HG00342', 'HG00345', 'HG00351', 'HG00358', 'HG00360', 'HG00366', 'HG00369', 'HG00371', 'HG00372', 'HG00375', 'HG00382', 'HG00403', 'HG00406', 'HG00409', 'HG00421', 'HG00436', 'HG00442', 'HG00445', 'HG00448', 'HG00451', 'HG00457', 'HG00463', 'HG00472', 'HG00475', 'HG00478', 'HG00500', 'HG00524', 'HG00530', 'HG00533', 'HG00536', 'HG00542', 'HG00553', 'HG00556', 'HG00559', 'HG00565', 'HG00580', 'HG00583', 'HG00589', 'HG00592', 'HG00595', 'HG00598', 'HG00607', 'HG00610', 'HG00613', 'HG00619', 'HG00622', 'HG00625', 'HG00628', 'HG00631', 'HG00634', 'HG00637', 'HG00640', 'HG00650', 'HG00653', 'HG00656', 'HG00662', 'HG00671', 'HG00674', 'HG00683', 'HG00689', 'HG00692', 'HG00698', 'HG00701', 'HG00704', 'HG00707', 'HG00728', 'HG00731', 'HG00736', 'HG00739', 'HG00742', 'HG00844', 'HG00881', 'HG00982', 'HG01028', 'HG01031', 'HG01047', 'HG01048', 'HG01051', 'HG01054', 'HG01060', 'HG01063', 'HG01066', 'HG01069', 'HG01072', 'HG01075', 'HG01079', 'HG01082', 'HG01085', 'HG01088', 'HG01094', 'HG01097', 'HG01101', 'HG01104', 'HG01107', 'HG01110', 'HG01112', 'HG01121', 'HG01124', 'HG01130', 'HG01133', 'HG01136',

'HG01139', 'HG01142', 'HG01148', 'HG01161', 'HG01164', 'HG01167', 'HG01170', 'HG01173', 'HG01176', 'HG01182', 'HG01187', 'HG01190', 'HG01197', 'HG01200', 'HG01204', 'HG01241', 'HG01247', 'HG01250', 'HG01253', 'HG01256', 'HG01259', 'HG01271', 'HG01277', 'HG01280', 'HG01286', 'HG01302', 'HG01305', 'HG01308', 'HG01311', 'HG01325', 'HG01334', 'HG01341', 'HG01344', 'HG01350', 'HG01353', 'HG01356', 'HG01359', 'HG01362', 'HG01365', 'HG01374', 'HG01377', 'HG01383', 'HG01389', 'HG01392', 'HG01395', 'HG01398', 'HG01402', 'HG01405', 'HG01412', 'HG01413', 'HG01431', 'HG01437', 'HG01440', 'HG01443', 'HG01455', 'HG01461', 'HG01464', 'HG01479', 'HG01485', 'HG01488', 'HG01491', 'HG01494', 'HG01497', 'HG01500', 'HG01503', 'HG01506', 'HG01509', 'HG01512', 'HG01515', 'HG01518', 'HG01521', 'HG01524', 'HG01527', 'HG01530', 'HG01536', 'HG01550', 'HG01556', 'HG01565', 'HG01571', 'HG01577', 'HG01583', 'HG01586', 'HG01589', 'HG01596', 'HG01603', 'HG01606', 'HG01608', 'HG01610', 'HG01615', 'HG01617', 'HG01619', 'HG01624', 'HG01625', 'HG01630', 'HG01631', 'HG01669', 'HG01672', 'HG01675', 'HG01678', 'HG01680', 'HG01682', 'HG01686', 'HG01694', 'HG01699', 'HG01700', 'HG01705', 'HG01708', 'HG01709', 'HG01747', 'HG01756', 'HG01761', 'HG01765', 'HG01767', 'HG01771', 'HG01775', 'HG01777', 'HG01781', 'HG01783', 'HG01785', 'HG01789', 'HG01791', 'HG01810', 'HG01811', 'HG01816', 'HG01840'

0.0, 0.0, 0.0, 0.0, 0.0, 0.7563025210084033, 0.0, 0.0, 0.0, 0.0, 0.0, 0.7563025210084033, 0.0, 0.8823529411764706, 0.7563025210084033, 0.0, 1.4663865546218486, 0.0, 0.0, 0.0, 0.0, 0.0, 0.0, 0.0, 0.0, 0.0, 0.0, 0.0, 0.0, 0.8823529411764706, 0.0, 0.0, 0.0, 0.0, 0.0, 0.0, 0.0, 0.7563025210084033, 0.0, 0.0, 0.0, 0.0, 0.0, 0.0, 0.0, 0.0, 0.0, 2.546218487394958, 0.0, 0.0, 0.0, 0.0, 0.8823529411764706, 0.0, 0.0, 0.0, 0.0, 0.0, 0.0, 0.0, 1.5924369747899159, 0.0, 0.0, 0.0, 0.0, 0.0, 0.0, 0.0, 0.0, 0.0, 0.0, 0.0, 0.0, 0.0, 0.0, 0.0, 0.0, 0.0, 0.0, 0.0, 0.0, 0.0, 0.0, 0.7563025210084033, 0.0, 0.0, 0.7100840336134453, 0.0, 0.7100840336134453, 0.7100840336134453, 0.7100840336134453, 0.7100840336134453, 0.7100840336134453, 0.0, 0.7100840336134453, 0.7100840336134453, 0.7100840336134453, 0.7100840336134453, 0.7100840336134453, 0.7100840336134453, 1.699579831932773, 0.0, 0.0, 0.7563025210084033,

0.7100840336134453, 4.496848739495799, 0.0, 0.7100840336134453, 0.0, 0.7100840336134453, 0.0, 0.7563025210084033, 0.0, 0.0, 0.7100840336134453, 0.7100840336134453, 0.7100840336134453, 0.0, 0.0, 0.0, 0.7100840336134453, 0.0, 0.0, 1.4663865546218486, 0.7100840336134453, 0.0, 0.7563025210084033, 0.7100840336134453, 0.0, 0.0, 0.0, 0.0, 0.7100840336134453, 0.0, 0.7563025210084033, 0.0, 0.0, 0.7100840336134453, 0.7563025210084033, 0.7100840336134453, 0.7100840336134453, 0.7563025210084033, 0.0, 0.7100840336134453, 0.0, 0.7100840336134453, 0.7100840336134453, 0.7100840336134453, 0.0, 0.7100840336134453, 0.7100840336134453, 0.0, 0.0, 0.0, 0.7100840336134453, 0.0, 0.0, 0.0, 0.8823529411764706, 0.0, 0.7563025210084033, 0.0, 0.0, 0.0, 0.7563025210084033, 0.0, 0.0, 0.0, 0.0, 0.0, 0.0, 0.0, 0.0, 0.0, 0.0, 0.0, 0.0, 0.0, 0.0, 0.0, 0.0, 0.0, 0.0, 0.0, 0.0, 0.8823529411764706, 0.7563025210084033, 0.0, 0.0, 0.0, 0.0, 0.7563025210084033, 0.0, 0.0, 0.0, 0.0, 0.953781512605042, 0.0, 0.7563025210084033, 0.0, 0.7563025210084033, 0.0, 0.0, 0.0, 0.0, 0.0, 0.0, 0.0, 0.0, 0.0, 0.9894957983193278, 0.0, 0.0, 0.0, 0.7563025210084033, 0.7100840336134453, 0.953781512605042, 0.0, 0.7563025210084033, 0.0, 0.0, 0.0, 0.0, 0.0, 0.0, 0.0, 0.0, 0.0, 0.7563025210084033, 0.0, 0.0, 0.0, 0.7563025210084033, 0.0, 0.0, 0.0, 0.7100840336134453, 0.0, 0.0, 0.0, 0.0, 0.0, 0.0, 0.0, 0.0, 0.0, 0.7563025210084033, 1.6638655462184873, 0.0, 0.0, 0.0, 0.0, 0.0, 0.0, 0.7100840336134453, 0.7100840336134453, 0.0, 0.7563025210084033, 0.0, 0.7100840336134453, 0.953781512605042, 0.7100840336134453, 0.7100840336134453, 0.0, 0.0, 0.7563025210084033, 0.7563025210084033, 0.0, 0.0, 0.0, 0.0, 0.7563025210084033, 0.7100840336134453, 0.7563025210084033, 0.7563025210084033, 0.0, 1.4663865546218486, 0.0, 0.7563025210084033, 0.0, 0.7563025210084033, 0.0, 0.7563025210084033, 0.0, 0.0, 0.7563025210084033, 0.0, 0.0, 0.0, 0.0, 0.0, 0.7563025210084033, 0.7100840336134453, 0.7100840336134453, 0.0, 0.0, 0.0, 0.0, 0.7100840336134453, 0.0, 0.7100840336134453

**(women): sample 1(HG00097), sample 2, sample 3, ... ,sample 300(HG01605) the total collective distance of sample 1(5.253151260504202), the total collective distance of sample 2, the total collective distance of sample 3, ..., the total collective distance of sample 300(5.253151260504202)**

'HG00097', 'HG00099', 'HG00100', 'HG00102', 'HG00104', 'HG00106', 'HG00110', 'HG00111', 'HG00118', 'HG00120', 'HG00121', 'HG00122', 'HG00123', 'HG00125', 'HG00127', 'HG00128', 'HG00130', 'HG00132', 'HG00133', 'HG00134', 'HG00135', 'HG00137', 'HG00146', 'HG00150', 'HG00154', 'HG00158', 'HG00171', 'HG00173', 'HG00174', 'HG00176', 'HG00177', 'HG00178', 'HG00179', 'HG00180', 'HG00231', 'HG00232', 'HG00233', 'HG00235', 'HG00236', 'HG00237', 'HG00238', 'HG00239', 'HG00240', 'HG00245', 'HG00249', 'HG00250', 'HG00253', 'HG00254', 'HG00255', 'HG00257', 'HG00258', 'HG00259', 'HG00261', 'HG00262', 'HG00263', 'HG00266', 'HG00268', 'HG00269', 'HG00270', 'HG00272', 'HG00274', 'HG00275', 'HG00276', 'HG00281', 'HG00282', 'HG00285', 'HG00288', 'HG00302', 'HG00304', 'HG00306', 'HG00309', 'HG00313', 'HG00315', 'HG00318', 'HG00319', 'HG00320', 'HG00323', 'HG00324', 'HG00326', 'HG00327', 'HG00328', 'HG00330', 'HG00331', 'HG00332', 'HG00334', 'HG00337', 'HG00339', 'HG00343', 'HG00344', 'HG00346', 'HG00349', 'HG00350', 'HG00353', 'HG00355', 'HG00356', 'HG00357', 'HG00359', 'HG00361', 'HG00362', 'HG00364', 'HG00365', 'HG00367', 'HG00368', 'HG00373', 'HG00376', 'HG00377', 'HG00378', 'HG00379', 'HG00380', 'HG00381', 'HG00383', 'HG00384', 'HG00404', 'HG00407', 'HG00410', 'HG00419', 'HG00422', 'HG00428', 'HG00437', 'HG00443', 'HG00446', 'HG00449', 'HG00452', 'HG00458', 'HG00464', 'HG00473', 'HG00476', 'HG00479', 'HG00513', 'HG00525', 'HG00531', 'HG00534', 'HG00537', 'HG00543', 'HG00551', 'HG00554', 'HG00557', 'HG00560', 'HG00566', 'HG00581', 'HG00584', 'HG00590', 'HG00593', 'HG00596', 'HG00599', 'HG00608', 'HG00611', 'HG00614', 'HG00620', 'HG00623', 'HG00626', 'HG00629', 'HG00632', 'HG00638', 'HG00641', 'HG00651', 'HG00654', 'HG00657', 'HG00663', 'HG00672', 'HG00675', 'HG00684', 'HG00690', 'HG00693', 'HG00699', 'HG00705', 'HG00708', 'HG00717', 'HG00729', 'HG00732', 'HG00734', 'HG00737', 'HG00740', 'HG00743', 'HG00759', 'HG00766', 'HG00851', 'HG00864', 'HG00867', 'HG00879', 'HG00956', 'HG00978',

'HG01029', 'HG01046', 'HG01049', 'HG01052', 'HG01055', 'HG01058', 'HG01061', 'HG01064', 'HG01067', 'HG01070', 'HG01073', 'HG01077', 'HG01080', 'HG01083', 'HG01086', 'HG01089', 'HG01092', 'HG01095', 'HG01098', 'HG01102', 'HG01105', 'HG01108', 'HG01111', 'HG01113', 'HG01119', 'HG01122', 'HG01125', 'HG01131', 'HG01134', 'HG01137', 'HG01140', 'HG01149', 'HG01162', 'HG01168', 'HG01171', 'HG01174', 'HG01177', 'HG01183', 'HG01188', 'HG01191', 'HG01198', 'HG01205', 'HG01242', 'HG01248', 'HG01251', 'HG01254', 'HG01257', 'HG01260', 'HG01269', 'HG01272', 'HG01275', 'HG01281', 'HG01284', 'HG01303', 'HG01312', 'HG01323', 'HG01326', 'HG01342', 'HG01345', 'HG01348', 'HG01351', 'HG01354', 'HG01357', 'HG01360', 'HG01363', 'HG01366', 'HG01369', 'HG01372', 'HG01375', 'HG01378', 'HG01384', 'HG01390', 'HG01393', 'HG01396', 'HG01403', 'HG01414', 'HG01432', 'HG01435', 'HG01438', 'HG01441', 'HG01444', 'HG01447', 'HG01456', 'HG01459', 'HG01462', 'HG01465', 'HG01468', 'HG01471', 'HG01474', 'HG01486', 'HG01489', 'HG01492', 'HG01495', 'HG01498', 'HG01501', 'HG01504', 'HG01507', 'HG01510', 'HG01513', 'HG01516', 'HG01519', 'HG01522', 'HG01525', 'HG01528', 'HG01531', 'HG01537', 'HG01551', 'HG01566', 'HG01572', 'HG01578', 'HG01593', 'HG01595', 'HG01597', 'HG01598', 'HG01599', 'HG01600', 'HG01602', 'HG01605’

5.253151260504202, 5.253151260504202, 5.253151260504202, 5.253151260504202, 5.253151260504202, 5.253151260504202, 5.253151260504202, 5.253151260504202, 5.253151260504202, 5.253151260504202, 5.253151260504202, 5.253151260504202, 5.253151260504202, 5.253151260504202, 5.253151260504202, 5.253151260504202, 5.253151260504202, 5.253151260504202, 5.253151260504202, 5.253151260504202, 5.253151260504202, 5.253151260504202, 5.253151260504202, 5.253151260504202, 5.253151260504202, 5.253151260504202, 5.253151260504202, 5.253151260504202, 5.253151260504202, 5.253151260504202, 5.253151260504202, 5.253151260504202, 5.253151260504202, 5.253151260504202, 5.253151260504202, 5.253151260504202, 5.253151260504202, 5.253151260504202, 5.253151260504202,

5.253151260504202, 5.253151260504202, 5.253151260504202,
5.253151260504202, 5.253151260504202, 5.253151260504202,
5.253151260504202, 5.253151260504202, 5.253151260504202,
5.253151260504202, 5.253151260504202, 5.253151260504202,
5.253151260504202, 5.253151260504202, 5.253151260504202,
5.253151260504202, 5.253151260504202, 5.253151260504202,
5.253151260504202, 5.253151260504202, 5.253151260504202,
5.253151260504202, 5.253151260504202, 5.253151260504202,
5.253151260504202, 5.253151260504202, 5.253151260504202,
5.253151260504202, 3.3203781512605044, 5.253151260504202,
5.253151260504202, 5.253151260504202, 5.253151260504202,
5.253151260504202, 4.281512605042017, 5.253151260504202,
5.253151260504202, 5.253151260504202, 5.253151260504202,
5.253151260504202, 5.253151260504202, 5.253151260504202,
5.253151260504202, 5.253151260504202, 5.253151260504202,
5.253151260504202, 5.253151260504202, 5.253151260504202,
5.253151260504202, 5.253151260504202, 5.253151260504202,
5.253151260504202, 5.253151260504202, 4.281512605042017,
4.496848739495799, 5.253151260504202, 5.253151260504202,
5.253151260504202, 5.253151260504202, 5.253151260504202,
5.253151260504202, 5.253151260504202, 5.253151260504202,
5.253151260504202, 5.253151260504202, 5.253151260504202,
4.496848739495799, 5.253151260504202, 5.253151260504202,
5.253151260504202, 5.253151260504202, 5.253151260504202,
5.253151260504202, 5.253151260504202, 5.253151260504202,
5.253151260504202, 5.253151260504202, 5.253151260504202,
5.253151260504202, 5.253151260504202, 5.253151260504202,
5.253151260504202, 5.253151260504202, 5.253151260504202,
5.253151260504202, 5.253151260504202, 5.253151260504202,
5.253151260504202, 5.253151260504202, 5.253151260504202,

5.253151260504202, 5.253151260504202, 5.253151260504202,
5.253151260504202, 5.253151260504202, 5.253151260504202,
5.253151260504202, 5.253151260504202, 5.253151260504202,
5.253151260504202, 5.253151260504202, 5.253151260504202,
5.253151260504202, 5.253151260504202, 5.253151260504202,
5.253151260504202, 5.253151260504202, 5.253151260504202,
5.253151260504202, 5.253151260504202, 5.253151260504202,
5.253151260504202, 5.253151260504202, 4.29936974789916,
5.253151260504202, 5.253151260504202, 5.253151260504202,
5.253151260504202, 5.253151260504202, 5.253151260504202,
5.253151260504202, 5.253151260504202, 5.253151260504202,
5.253151260504202, 5.253151260504202, 5.253151260504202,
5.253151260504202, 5.253151260504202, 5.253151260504202,
5.253151260504202, 5.253151260504202, 4.496848739495799,
5.253151260504202, 5.253151260504202, 5.253151260504202,
5.253151260504202, 5.253151260504202, 5.253151260504202,
5.253151260504202, 5.253151260504202, 5.253151260504202,
5.253151260504202, 5.253151260504202, 5.253151260504202,
5.253151260504202, 5.253151260504202, 5.253151260504202,
5.253151260504202, 5.253151260504202, 5.253151260504202,
5.253151260504202, 5.253151260504202, 5.253151260504202,
5.253151260504202, 5.253151260504202, 5.253151260504202,
5.253151260504202, 5.253151260504202, 5.253151260504202,
5.253151260504202, 5.253151260504202, 5.253151260504202,
5.253151260504202, 5.253151260504202, 5.253151260504202,
5.253151260504202, 5.253151260504202, 5.253151260504202,
5.253151260504202, 5.253151260504202, 5.253151260504202,
5.253151260504202, 5.253151260504202, 5.253151260504202,
5.253151260504202, 5.253151260504202, 5.253151260504202,
5.253151260504202, 5.253151260504202, 5.253151260504202,

5.253151260504202, 5.253151260504202, 5.253151260504202,
5.253151260504202, 5.253151260504202, 5.253151260504202,
5.253151260504202, 5.253151260504202, 5.253151260504202,
5.253151260504202, 5.253151260504202, 5.253151260504202,
5.253151260504202, 5.253151260504202, 5.253151260504202,
5.253151260504202, 5.253151260504202, 5.253151260504202,
5.253151260504202, 5.253151260504202, 5.253151260504202,
5.253151260504202, 5.253151260504202, 5.253151260504202,
5.253151260504202, 5.253151260504202, 5.253151260504202,
5.253151260504202, 5.253151260504202, 5.253151260504202,
5.253151260504202, 5.253151260504202, 5.253151260504202,
5.253151260504202, 5.253151260504202, 5.253151260504202,
5.253151260504202, 5.253151260504202, 5.253151260504202,
5.253151260504202, 5.253151260504202, 5.253151260504202,
5.253151260504202, 5.253151260504202, 5.253151260504202,
5.253151260504202, 5.253151260504202, 5.253151260504202,
5.253151260504202, 5.253151260504202, 5.253151260504202,
5.253151260504202, 5.253151260504202, 5.253151260504202,
4.496848739495799, 5.253151260504202, 5.253151260504202,
5.253151260504202, 5.253151260504202, 5.253151260504202,
5.253151260504202, 5.253151260504202, 5.253151260504202,
5.253151260504202, 5.253151260504202, 5.253151260504202,
5.253151260504202, 5.253151260504202, 4.281512605042017,
5.253151260504202, 5.253151260504202, 5.253151260504202,
5.253151260504202, 5.253151260504202, 5.253151260504202,
5.253151260504202, 5.253151260504202, 5.253151260504202,
5.253151260504202, 5.253151260504202, 5.253151260504202

**1.4. The total collective distances for 51 SNPs(0.07 ≤ $|D_{iXY}|$ ≤ 0.21) (men) : same sample order above**

27.366596638655462, 26.802521008403364, 25.13235294117647, 25.13340336134454, 25.86764705882353, 25.211134453781515, 25.051470588235297, 24.473739495798323, 28.6281512605042, 29.613445378151262, 26.347689075630257, 25.188025210084035, 28.102941176470587, 31.796218487394956, 28.789915966386555, 27.58298319327731, 29.29516806722689, 27.390756302521012, 27.70168067226891, 26.192226890756306, 28.772058823529413, 28.983193277310924, 27.69012605042017, 29.23004201680672, 29.316176470588236, 30.015756302521005, 28.025210084033613, 28.694327731092436, 29.807773109243698, 29.17752100840336, 28.108193277310924, 27.219537815126053, 23.926470588235297, 27.825630252100844, 27.21638655462185, 27.83718487394958, 29.235294117647058, 29.201680672268907, 30.301470588235297, 29.057773109243698, 25.873949579831937, 26.760504201680675, 28.403361344537814, 26.46218487394958, 27.51575630252101, 25.595588235294116, 29.38655462184874, 32.15966386554622, 27.460084033613445, 24.017857142857146, 25.320378151260506, 29.3109243697479, 29.43487394957983, 26.218487394957982, 29.130252100840337, 26.313025210084035, 25.415966386554622, 27.34873949579832, 29.342436974789916, 27.608193277310924, 29.91176470588235, 29.709033613445378, 26.08403361344538, 26.298319327731093, 28.450630252100844, 28.910714285714285, 27.285714285714285, 23.934873949579835, 29.23949579831933, 29.095588235294116, 30.827731092436977, 24.953781512605044, 29.11764705882353, 31.051470588235293, 26.429621848739497, 27.403361344537817, 26.377100840336134, 28.51365546218487, 27.69327731092437, 28.369747899159663, 29.769957983193276, 27.922268907563023, 28.792016806722692, 30.04516806722689,

30.449579831932773, 29.063025210084035, 28.574579831932773,
29.820378151260506, 27.03046218487395, 30.764705882352942,
27.75735294117647, 30.431722689075627, 26.98109243697479,
26.78256302521008, 25.295168067226893, 27.646008403361343,
26.477941176470587, 27.860294117647058, 28.870798319327733,
25.975840336134457, 25.230042016806724, 28.463235294117645,
25.15966386554622, 26.71743697478992, 30.28361344537815,
30.346638655462186, 24.563025210084035, 29.019957983193276,
29.468487394957982, 28.22373949579832, 27.682773109243698,
28.391806722689076, 23.82247899159664, 24.871848739495803,
27.05672268907563, 25.750000000000004, 28.42436974789916,
26.73109243697479, 28.464285714285715, 32.66386554621849,
27.49264705882353, 29.294117647058822, 29.16806722689076,
23.92226890756303, 28.69012605042017, 24.259453781512608,
28.174369747899163, 30.217436974789916, 30.080882352941174,
28.404411764705884, 28.60609243697479, 31.660714285714285,
29.53046218487395, 29.563025210084035, 28.77310924369748,
26.20168067226891, 30.94747899159664, 27.56617647058824,
28.76575630252101, 30.08823529411765, 26.866596638655462,
28.165966386554622, 29.59873949579832, 23.460084033613445,
28.719537815126053, 27.274159663865547, 25.25735294117647,
26.528361344537817, 29.797268907563026, 29.988445378151262,
27.997899159663866, 27.84138655462185, 26.836134453781515,
25.222689075630257, 25.488445378151262, 29.62079831932773,
27.733193277310924, 27.563025210084035, 29.80672268907563,
27.93487394957983, 27.318277310924373, 26.864495798319332,
29.547268907563026, 29.02941176470588, 26.743697478991596,
27.620798319327733, 30.169117647058826, 23.853991596638657,
26.923319327731093, 23.237394957983195, 32.27205882352941,
29.284663865546218, 30.15126050420168, 29.52310924369748,

29.410714285714285, 29.023109243697483, 29.040966386554622,
28.63865546218487, 24.46323529411765, 22.438025210084035,
29.04516806722689, 23.550420168067227, 28.798319327731093,
27.490546218487395, 28.691176470588236, 31.28571428571429,
29.53781512605042, 27.95483193277311, 30.493697478991596,
29.338235294117645, 29.061974789915965, 30.71218487394958,
27.78676470588235, 28.10504201680672, 28.485294117647058,
26.41281512605042, 28.238445378151262, 27.58613445378151,
28.14495798319328, 26.492647058823533, 27.16386554621849,
23.903361344537817, 26.20168067226891, 29.671218487394956,
24.007352941176475, 27.532563025210088, 27.73424369747899,
27.16281512605042, 27.990546218487395, 23.78886554621849,
24.585084033613448, 26.475840336134453, 29.289915966386555,
30.149159663865547, 30.220588235294116, 28.26890756302521,
28.56197478991597, 30.261554621848738, 29.45483193277311,
22.85924369747899, 25.949579831932773, 30.19642857142857,
29.084033613445378, 31.451680672268907, 30.7436974789916,
29.40441176470588, 28.699579831932773, 27.11764705882353,
29.89075630252101, 29.23004201680672, 26.45168067226891,
27.727941176470587, 23.415966386554622, 29.194327731092436,
28.554621848739497, 24.033613445378155, 27.410714285714285,
25.982142857142858, 22.711134453781515, 26.467436974789916,
26.71113445378151, 29.304621848739494, 24.978991596638654,
24.457983193277315, 23.76470588235294, 25.130252100840337,
27.46638655462185, 24.03571428571429, 22.790966386554626,
28.392857142857142, 23.14075630252101, 26.617647058823533,
22.990546218487395, 27.99264705882353, 25.7436974789916,
25.718487394957986, 19.671218487394963, 23.287815126050422,
27.282563025210084, 28.067226890756302, 26.988445378151262,
26.96953781512605, 26.648109243697483, 25.282563025210088,

25.806722689075627, 27.52310924369748, 27.990546218487395,
26.911764705882355, 31.057773109243698, 27.804621848739494,
29.490546218487395, 23.83403361344538, 24.2468487394958,
25.245798319327733, 28.258403361344538, 28.327731092436977,
27.13865546218487, 27.491596638655466, 25.858193277310928,
24.7436974789916, 28.45378151260504, 29.220588235294116,
27.94012605042017, 23.620798319327733, 28.534663865546218,
26.971638655462183, 25.872899159663866, 30.801470588235297,
29.23109243697479, 25.07142857142857, 29.10924369747899,
27.387605042016805, 30.463235294117645, 23.648109243697483,
23.26575630252101, 29.60609243697479, 28.25735294117647,
27.021008403361346, 26.693277310924373, 26.79936974789916

**(women) : same sample order above**

31.338235294117645, 30.843487394957982, 31.15126050420168,
30.80987394957983, 33.06512605042017, 31.765756302521005,
30.680672268907564, 32.753151260504204, 30.897058823529413,
33.036764705882355, 34.57668067226891, 33.305672268907564,
32.596638655462186, 33.33718487394958, 33.523109243697476,
33.43172268907563, 30.27626050420168, 32.89285714285714,
34.14705882352941, 35.59348739495798, 30.44327731092437,
32.463235294117645, 33.44012605042017, 32.46533613445378,
32.55042016806723, 32.71638655462185, 35.515756302521005,
32.45483193277311, 32.148109243697476, 31.070378151260506,
32.51575630252101, 33.739495798319325, 33.49159663865546,
30.179621848739494, 34.38865546218487, 32.73529411764706,
31.264705882352942, 31.560924369747898, 33.96638655462185,
31.551470588235293, 33.32142857142857, 30.102941176470587,
31.5640756302521, 33.40126050420168, 32.58718487394958,
33.720588235294116, 35.174369747899156, 31.502100840336134,

31.65861344537815, 34.061974789915965, 34.6922268907563,
33.25735294117647, 31.758403361344534, 32.116596638655466,
32.60609243697479, 32.528361344537814, 33.113445378151255,
32.417016806722685, 32.96848739495798, 30.850840336134453,
32.419117647058826, 33.627100840336134, 33.589285714285715,
32.167016806722685, 31.188025210084035, 30.956932773109244,
32.54726890756302, 32.78361344537815, 33.08298319327731,
34.51470588235294, 33.213235294117645, 34.26890756302521,
33.34873949579832, 32.38865546218487, 32.62394957983193,
31.023109243697476, 32.13655462184874, 31.254201680672267,
31.293067226890756, 32.46638655462185, 31.827731092436974,
34.42016806722689, 33.85924369747899, 31.92016806722689,
31.68382352941176, 34.44117647058823, 30.876050420168067,
33.829831932773104, 31.37394957983193, 34.61029411764706,
34.20168067226891, 34.213235294117645, 34.0063025210084,
30.963235294117645, 33.26995798319328, 34.3781512605042,
36.15756302521008, 33.73109243697479, 34.39600840336134,
32.121848739495796, 31.544117647058826, 33.383403361344534,
33.161764705882355, 33.09768907563025, 30.963235294117645,
32.496848739495796, 33.286764705882355, 32.03991596638655,
32.45063025210084, 33.214285714285715, 32.85504201680672,
30.558823529411764, 32.469537815126046, 31.48424369747899,
33.0126050420168, 32.96218487394958, 34.80987394957983,
34.42226890756302, 34.08718487394958, 33.80567226890756,
32.220588235294116, 33.002100840336134, 32.89075630252101,
31.767857142857142, 32.12605042016806, 33.37920168067227,
33.94747899159664, 32.65861344537815, 32.240546218487395,
33.0577731092437, 32.41281512605042, 34.32563025210084,
31.834033613445378, 32.59033613445378, 31.42016806722689,
33.73739495798319, 30.86764705882353, 35.09978991596638,

32.365546218487395, 33.569327731092436, 33.74264705882353,
32.14285714285714, 33.384453781512605, 33.0, 33.35714285714286,
33.8172268907563, 34.0672268907563, 32.6376050420168,
31.54621848739496, 33.22478991596638, 32.18172268907563,
32.8172268907563, 33.509453781512605, 32.67121848739495,
32.15021008403362, 32.20378151260504, 31.527310924369747,
32.48529411764706, 31.817226890756302, 34.75525210084033,
32.65651260504202, 33.47899159663865, 32.5735294117647,
32.15651260504202, 33.0735294117647, 34.174369747899156,
33.252100840336134, 31.971638655462186, 33.66596638655462,
31.4968487394958, 32.95903361344538, 31.48109243697479,
34.28886554621849, 31.807773109243698, 32.98529411764706,
32.52415966386555, 34.09978991596638, 32.469537815126046,
34.45798319327731, 33.903361344537814, 31.07142857142857,
33.17647058823529, 31.619747899159663, 32.944327731092436,
34.497899159663866, 30.811974789915965, 33.41386554621849,
34.20483193277311, 32.02731092436974, 31.810924369747898,
33.38655462184874, 32.022058823529406, 32.15021008403361,
32.36869747899159, 31.113445378151262, 31.092436974789916,
33.02415966386555, 30.630252100840337, 30.25, 31.717436974789916,
30.11764705882353, 33.88970588235294, 31.871848739495796,
34.290966386554615, 32.46008403361344, 31.744747899159663,
32.36134453781512, 33.26575630252101, 30.352941176470587,
32.26260504201681, 34.10084033613445, 33.0577731092437,
33.76365546218487, 33.54936974789916, 32.08298319327731,
29.9359243697479, 30.464285714285715, 32.51995798319328,
30.005252100840337, 32.13655462184874, 33.23109243697479,
31.722689075630257, 34.35504201680672, 33.213235294117645,
32.21113445378151, 31.57878151260504, 33.12394957983193,
32.470588235294116, 30.626050420168067, 31.839285714285715,

31.66281512605042, 33.15861344537815, 32.338235294117645,
31.195378151260506, 32.6922268907563, 32.71218487394958,
33.990546218487395, 33.14285714285714, 30.926470588235293,
33.26470588235294, 30.956932773109244, 33.36344537815126,
33.580882352941174, 32.570378151260506, 32.410714285714285,
33.14285714285714, 32.266806722689076, 34.36239495798319,
34.1890756302521, 32.01470588235294, 30.607142857142854,
30.876050420168063, 33.471638655462186, 32.61764705882353,
31.67436974789916, 33.55987394957983, 33.29516806722689,
33.22794117647059, 31.189075630252102, 34.069327731092436,
32.13865546218487, 33.04516806722689, 32.87079831932773,
34.238445378151255, 32.83508403361344, 33.36239495798319,
32.99579831932773, 31.399159663865547, 30.630252100840337,
34.561974789915965, 34.51575630252101, 33.94957983193277,
29.376050420168067, 29.863445378151262, 32.00525210084034,
33.444327731092436, 30.85609243697479, 32.19852941176471,
30.27626050420168, 33.41281512605042, 31.474789915966383,
32.22794117647059, 31.26995798319328, 30.827731092436974,
31.62079831932773, 32.61869747899159, 31.608193277310924,
34.49579831932773, 33.77731092436974, 33.44957983193277,
34.419117647058826, 32.08613445378151, 31.648109243697476,
33.628151260504204, 33.71428571428571, 32.97584033613445,
33.930672268907564, 33.41491596638655, 32.721638655462186,
33.1376050420168
